# PLK-1 regulates MEX-1 polarization in the *C. elegans* zygote

**DOI:** 10.1101/2024.07.26.605193

**Authors:** Amelia J. Kim, Stephanie I. Miller, Elora C. Greiner, Arminja N. Kettenbach, Erik E. Griffin

## Abstract

The one-cell *C. elegans* embryo undergoes an asymmetric cell division during which germline factors such as the RNA-binding proteins POS-1 and MEX-1 segregate to the posterior cytoplasm, leading to their asymmetric inheritance to the posterior germline daughter cell. Previous studies found that the RNA-binding protein MEX-5 recruits polo-like kinase PLK-1 to the anterior cytoplasm where PLK-1 inhibits the retention of its substrate POS-1, leading to POS-1 segregation to the posterior. In this study, we tested whether PLK-1 similarly regulates MEX-1 polarization. We find that both the retention of MEX-1 in the anterior and the segregation of MEX-1 to the posterior depend on PLK kinase activity and on the interaction between MEX-5 and PLK-1. Human PLK1 directly phosphorylates recombinant MEX-1 on 9 predicted PLK-1 sites in vitro, four of which were identified in previous phosphoproteomic analysis of *C. elegans* embryos. The introduction of alanine substitutions at these four PLK-1 phosphorylation sites (MEX-1(4A)) significantly weakened the inhibition of MEX-1 retention in the anterior, thereby weakening MEX-1 segregation to the posterior. In contrast, mutation of a predicted CDK1 phosphorylation site had no effect on MEX-1 retention or on MEX-1 segregation. MEX-1(4A) mutants are viable and fertile but display significant sterility and fecundity defects at elevated temperatures. Taken together with our previous findings, these findings suggest PLK-1 phosphorylation drives both MEX-1 and POS-1 polarization during the asymmetric division of the zygote.

## INTRODUCTION

Asymmetric cell divisions generate daughter cells that differ in size, fate and/or function and are important for cell diversification during embryonic development (Li, 2013). During many asymmetric divisions, the polarization of the mother cell leads to the segregation of fate determinants to one pole, resulting in their asymmetric inheritance to one daughter cell at cell division (Sunchu and Cabernard, 2020). Because the polarization of fate determinants must be coordinated with the progression to cytokinesis, characterizing the interplay between cell cycle and cell polarity mechanisms is central to understanding how cells divide asymmetrically.

The asymmetric division of the *C. elegans* embryo gives rise to a somatic daughter cell and a germline daughter cell. As the newly fertilized embryo progresses through meiosis, a collection of maternally deposited RNA-binding proteins remain symmetrically distributed in the cytoplasm. Following the completion of meiosis, the embryo begins to polarize along the anterior/posterior axis, leading to the segregation of somatic factors to the anterior cytoplasm and germline factors to the posterior cytoplasm (Griffin, 2015; Lang and Munro, 2017; Peglion and Goehring, 2019). When the zygote divides ∼20 minutes after the completion of meiosis, these cytoplasmic factors are inherited asymmetrically, giving rise to an anterior somatic daughter AB and a posterior germline daughter cell P1. Three subsequent asymmetric divisions in the descendants of P1 each generate a somatic and a germline daughter (Rose and Gonczy, 2014; Wang and Seydoux, 2013). As a result of these asymmetries, the translation of maternal mRNAs encoding key signaling molecules and transcription factors is restricted to a subset of cells in the early embryo (Hwang and Rose, 2010).

The polarization of the zygote is orchestrated by the PAR proteins, which form distinct anterior and posterior PAR domains at the cell cortex (Lang and Munro, 2017). The posterior PAR kinase PAR-1 drives the redistribution of the redundant RNA-binding proteins MEX-5 and MEX-6 (MEX-5/6 hereafter) to the anterior cytoplasm by inhibiting their retention in the posterior cytoplasm (Daniels et al., 2010; Griffin et al., 2011; Schubert et al., 2000; Tenlen et al., 2008; Wu et al., 2018). As the embryo initiates polarization, MBK-2 kinase is activated and phosphorylates a polo-docking site on MEX-5/6, leading to the association of the polo-like kinase PLK-1 with MEX-5/6 (Nishi et al., 2008). This association results in the formation of an anterior-rich PLK-1 gradient that mirrors the MEX-5/6 gradient (Barbieri et al., 2022; Budirahardja et al., 2008; Chase et al., 2000; Nishi et al., 2008; Rivers et al., 2008).

As the MEX-5/6 and PLK-1 gradients form, the tandem CCCH zinc finger RNA-binding proteins POS-1, MEX-1 and PIE-1 segregate to the posterior cytoplasm, leading to their preferential inheritance by P1 (Guedes and Priess, 1997; Mello et al., 1996; Tabara et al., 1999). The low levels of POS-1, MEX-1 and PIE-1 inherited by AB are degraded in somatic cells, reinforcing their enrichment in the germline lineage (DeRenzo et al., 2003). During polarization, POS-1, MEX-1 and PIE-1 are retained in slow-diffusing complexes in the posterior cytoplasm, which underlies their segregation to the posterior (Daniels et al., 2009; Han et al., 2018; Wu et al., 2018; Wu et al., 2015). POS-1 and PIE-1 retention depend on their ability to bind RNA, suggesting they are retained in slow-diffusing RNA complexes in the posterior (Han et al., 2018; Wu et al., 2018). PLK-1 phosphorylation inhibits POS-1 retention in the anterior (Han et al., 2018). Mutation of the polo-docking site on MEX-5 renders PLK-1 incapable of inhibiting of POS-1 retention, suggesting it is PLK-1 in complex with MEX-5/6 that acts on POS-1 (Han et al., 2018). MEX-5/6 also control the segregation of PIE-1 and MEX-1 by inhibiting their retention in the anterior (Schubert et al., 2000; Wu et al., 2018; Wu et al., 2015), but the underlying mechanisms have not been established.

This study focuses on the mechanisms underlying MEX-1 segregation. MEX-1 is an essential, maternally contributed protein that contributes to the lineage-restricted localization of SKN-1, ZIF-1 and MOM-2 (Bowerman et al., 1993; Guedes and Priess, 1997; Mello et al., 1992; Oldenbroek et al., 2012; Oldenbroek et al., 2013). MEX-1 is also required for the segregation of P granules to the germline cell during the asymmetric divisions of P1 and its descendants (Mello et al., 1992; Schnabel et al., 1996). In *mex-1* mutant embryos, the fates of both somatic and germline founder cells are altered and embryos arrest at morphogenesis (Mello et al., 1992; Schnabel et al., 1996). Like PIE-1 and POS-1, MEX-1 is both associated with P granules in the posterior and forms a posterior-rich gradient in the cytoplasm surrounding P granules (Guedes and Priess, 1997). Here, we provide evidence that MEX-1 is a PLK-1 substrate and that PLK-1 phosphorylation inhibits MEX-1 retention in the anterior cytoplasm, leading to MEX-1 segregation to the posterior cytoplasm. These findings suggest that similar mechanisms underlie the polarization of POS-1 and MEX-1 during the asymmetric division of the zygote.

## RESULTS

### PLK-1 inhibits MEX-1 retention in the anterior

To begin to dissect the mechanisms that control MEX-1 segregation, we first characterized the dynamics of endogenously tagged MEX-1::GFP (Gauvin et al., 2018). MEX-1::GFP localizes to P granules and is enriched in the posterior cytoplasm outside of P granules (Guedes and Priess, 1997), forming a roughly 2-fold posterior-rich gradient (Figure 1A and 1B). We used FRAP (Fluorescence Recovery After Photobleaching) assays to monitor MEX-1::GFP mobility in the anterior and posterior cytoplasm at nuclear envelope breakdown (NEBD). The FRAP recovery of MEX-1::GFP is slower in the posterior than in the anterior cytoplasm, indicating that MEX-1::GFP is preferentially retained in the posterior (Figure 1C). The anterior-rich RNA-binding proteins MEX-5/6 are required for MEX-1 segregation (Schubert et al., 2000). In *mex-5/6(RNAi)* embryos, MEX-1::GFP is symmetrically distributed and its mobility in both the anterior and posterior cytoplasm is similar to its mobility in the posterior of wildtype embryos. Therefore, MEX-5/6 inhibits MEX-1::GFP retention in the anterior, driving its accumulation in the posterior (Figure 1A – 1C). These data are consistent with a previous study that used fluorescence correlation spectroscopy to characterize the dynamics of transgenic GFP::MEX-1 in the zygote (Wu et al., 2015).

**Figure 1.**
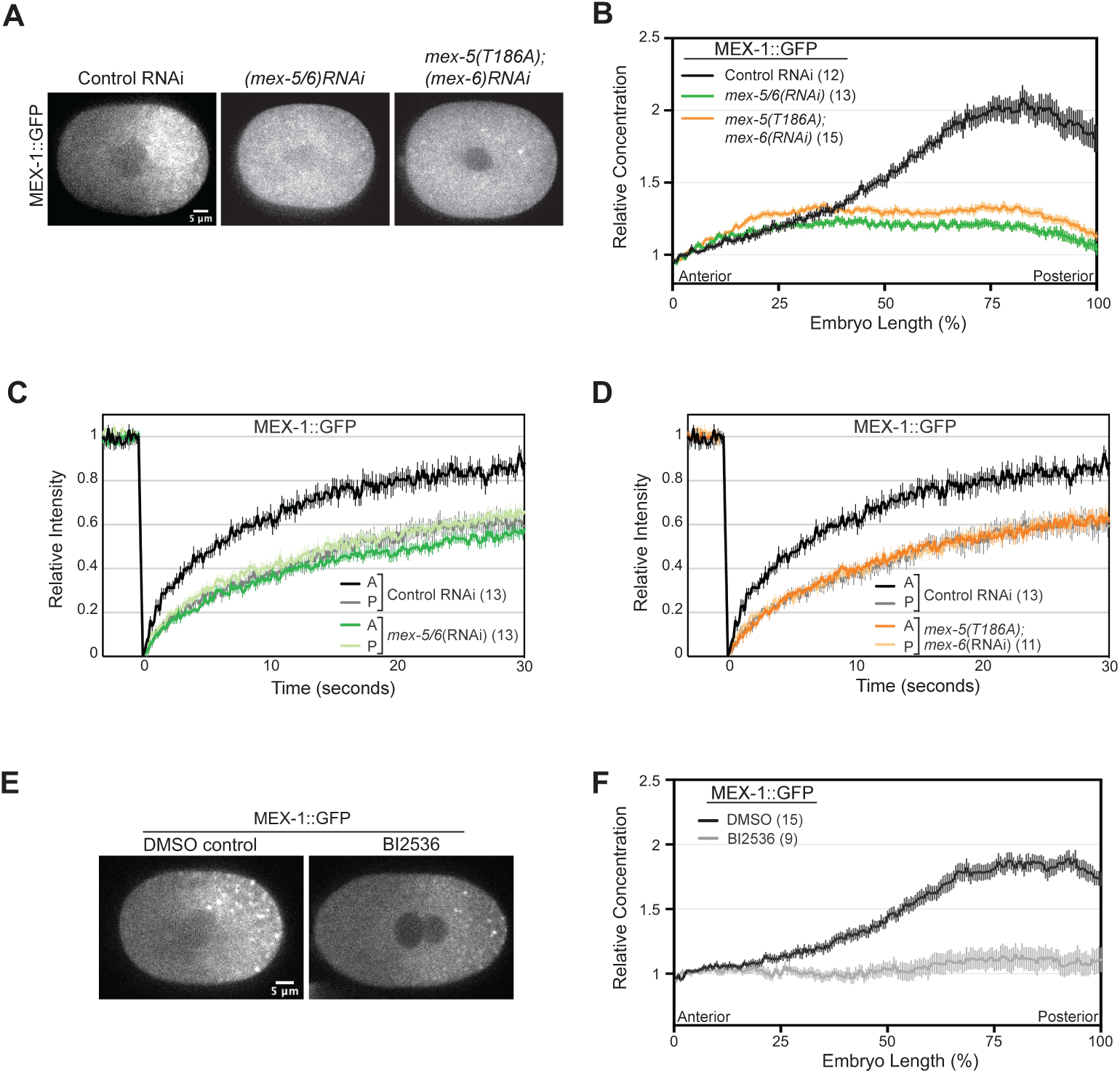
PLK-1 and MEX-5/6 control MEX-1 segregation in the *C. elegans* zygote. **A.** Fluorescence micrographs of MEX-1::GFP zygotes at nuclear envelope breakdown (NEBD). **B.** Average MEX-1::GFP fluorescence intensity along the anterior/posterior axis at NEBD. Intensities from the indicated number of embryos were normalized to the anterior end and averaged. **C, D.** Normalized FRAP (fluorescence recovery after photobleaching) curves following photobleaching in the anterior (labeled A) or posterior (labeled P) cytoplasm of zygotes at NEBD. The FRAP curves for the indicated number of embryos were normalized and averaged. **E.** Fluorescence micrographs of permeabilized zygotes treated with DMSO or BI2536, which inhibits polo-like kinases. **F.** Average MEX-1::GFP fluorescence intensity along the anterior/posterior axis following treatment with DMSO or BI2536. Intensities from the indicated number of embryos were were normalized to the anterior end and averaged. For E and F, *perm-1(RNAi)* was used to permeabilize embryos. For all graphs, error bars indicate SEM and the number of embryos analyzed is in parentheses.

The interaction between MEX-5/6 and PLK-1 kinase leads to the enrichment of PLK-1 in the anterior cytoplasm of the polarized zygote (Barbieri et al., 2022; Nishi et al., 2008). To test whether the interaction between MEX-5 and PLK-1 is required for MEX-1 segregation, we analyzed MEX-1::GFP dynamics in *mex-5(T186A);mex-6(RNAi)* embryos. T186A disrupts the interaction between PLK-1 and MEX-5 by preventing phosphorylation on the MEX-5 polo-docking site by MBK-2 kinase (Nishi et al., 2008). MEX-5 and MEX-6 are partially redundant (Schubert et al., 2000) and MEX-1::GFP segregates to the posterior in both *mex-6(RNAi)* and *mex-5(T186A)* embryos (Figure S1A). However, in *mex-5(T186A);mex-6(RNAi)* embryos, MEX-1::GFP fails to segregate and displays slow mobility in both the anterior and posterior cytoplasm, similar to *mex-5/6(RNAi)* embryos (Figure 1A, 1B and 1D). To test whether PLK kinase activity is required for MEX-1 segregation, we treated *perm-1(RNAi)* permeabilized one-cell embryos (Carvalho et al., 2011) with BI2536, which inhibits PLK kinase (Steegmaier et al., 2007). We find that MEX-1::GFP is symmetrically distributed in BI2536-treated embryos (Figure 1E). We conclude that PLK-1 acts in association with MEX-5/6 to regulate MEX-1 segregation.

### MEX-1 is a PLK-1 substrate

A previous phospho-proteomics analysis of *C. elegans* embryos identified five phosphorylation sites on MEX-1 (Offenburger et al., 2017). Four of these sites (S98, T235, S240, S248) are within the PLK consensus motif (D/E)-X-(S/T)-Φ-X-(D/E) (Elia et al., 2003; Nakajima et al., 2003) (Figure 2A). The fifth residue, S227, is within a CDK consensus motif (S/T-P-X-K/R) (Moreno and Nurse, 1990; Nigg, 1993; Songyang et al., 1994). Consistent with the possibility that PLK-1 might phosphorylate MEX-1, MEX-1 was detected in a PLK-1 proximity-labeling interactome study (Holzer et al., 2022). To test whether PLK1 can directly phosphorylate MEX-1, we performed *in vitro* kinase assays using human PLK1. Because we were unable to purify sufficient full-length MBP:MEX-1:6xHis following bacterial expression, we used an N-terminal fragment (aa1-299) that contains the 5 phosphorylation sites detected in embryos. PLK1 phosphorylated MBP:MEX-1(aa1-299):6xHis but not MBP *in vitro* (Figure 2A-C). Mutation of the 5 residues detected *in vivo* to alanine (“5A” mutant hereafter) significantly reduced *in vitro* phosphorylation of MEX-1 by PLK1, suggesting that the primary phosphorylation sites *in vitro* are among these residues. Phosphorylation was mapped to 9 sites by phospho-MS, including 6 of 7 predicted to be PLK-1 phosphorylation sites by the prediction program GPS-POLO 3.0 (Liu et al., 2013), or the Eukaryotic Linear Motif (ELM) (Kumar et al., 2022) and not including the predicted CDK site S227 (Figure 2A).

**Figure 2.**
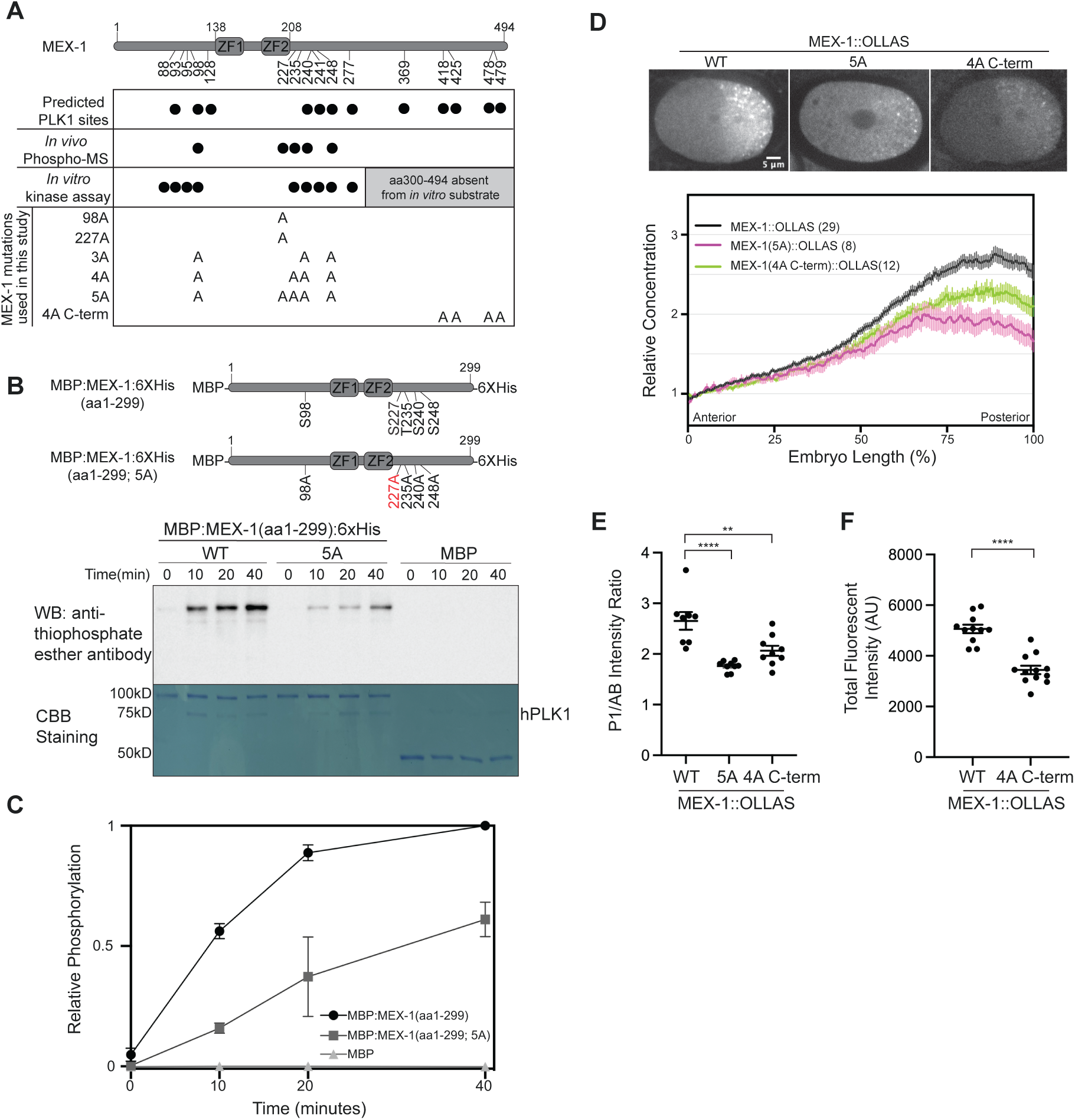
MEX-1 is a PLK1 substrate. **A.** Schematic of MEX-1 including the position of the RNA-binding zinc finger domains ZF1 and ZF2. Black circles in the panel below shows the position of predicted PLK1 phos-phorylation sites and phosphorylated residues detected by phospho-mass spectrometry of embryo lysates (*in vivo*; Offenburger 2017) or following *in vitro* kinase assays (as in panel B). The position of alanine substitutions in MEX-1 alleles used in this study are shown. Note that alleles in the N2, MEX-1::OLLAS and MEX-1::GFP backgrounds were made independently. **B.** *In vitro* kinase assay with hPLK1 and the indicated substrates. Top panel: Phosphorylation was detected by western blot using an anti-thiophosphate ester antibody, which recognizes alkylated ATP-γS. Bottom panel: total protein was detected by Coommassie Brilliant Blue (CBB) staining. The position of human hPLK1 is indicated to the right and the position of molecular weight markers (not shown) is indicated to the left. Schematic of the recombinant MEX-1 constructs used are shown above. **C.** Quantification hPLK1 phosphorylation of over time. Three replicates were normalized to the background (equals 0) the final MBP:MEX-1(aa1-299):6XHis value (equals 1) within each experiment and averaged. **D.** Top: Fluorescence micrographs of one-cell embryos immunostained using an anti-OLLAS antibody. Bottom: Average fluorescence intensity along the anterior/posterior axis of immunostained embryos. Intensities from the indicated number of embryos were normalized to the anterior end and averaged. **E.** Ratio of P1/AB fluorescence intensity in 2-cell embryos stained for MEX-1::OLLAS. Each dot indicates an individual embryo. The mean and SEM are shown. **F.** Total fluorescence intensity of immunostained MEX-1::OLLAS and MEX-1(4A C-term)::OLLAS zygotes. In this and subsequent figures, **** = p < 0.0001; *** = p < 0.001*, ** = p < 0.01, * = p < 0.05., n.s. = not significant.

We next introduced alanine substitutions at the 5 phosphorylation sites detected by Offenburger et al. (2017) using CRISPR/Cas9 gene editing. These mutations were first introduced into a MEX-1::OLLAS strain so that we could characterize MEX-1 localization by immunofluorescence using an OLLAS antibody. Strikingly, the segregation of MEX-1(5A)::OLLAS to the posterior cytoplasm before cell division and to the P1 daughter cell after cell division was significantly weaker than wild-type MEX-1::OLLAS (Figure 2D and 2E). Mutation of four predicted PLK-1 phosphorylation sites at the C-terminus (MEX-1(4A-C-term)::OLLAS) weakened MEX-1 segregation to the posterior before cell division and the enrichment in P1 after cell division (Figure 2D, 2E), indicating these residues also contribute to MEX-1 segregation. However, because the staining intensity of MEX-1(4A C-term)::OLLAS was lower than MEX-1::OLLAS (Figure 2F), because previous phosphoproteomic studies did not detect phosphorylation at these residues and because the MEX-1 fragment we used in our *in vitro* kinase assay did not include these residues, we did not consider them in our subsequent analysis.

### MEX-1 phosphorylation inhibits its retention in the anterior

To characterize the role of MEX-1 phosphorylation in the regulation of MEX-1 dynamics, we generated a series of alanine-substitution MEX-1::GFP alleles (Figure 2A). Similar to our findings with MEX-1(5A)::OLLAS, the segregation of MEX-1(5A)::GFP in the one-cell embryo was significantly weaker than MEX-1::GFP (Figure 3A and 3B). In the course of making MEX-1(5A)::GFP, we isolated strains with alanine substitutions at one (S98), three (S98, S240, S248; “3A”) or four (S98, T235, S240, S248; “4A”) PLK-1 phosphorylation sites, which caused progressively weaker MEX-1 segregation (Figure 3A and 3B). In contrast, mutation of S227, which lies within a CDK consensus sequence, did not disrupt MEX-1::GFP segregation (Figure 3A and 3B). We again used FRAP to monitor MEX-1::GFP retention in the anterior and posterior cytoplasm. We observed a significant increase in the retention of both MEX-5(5A)::GFP and MEX-1(4A)::GFP in the anterior cytoplasm relative to wildtype MEX-1::GFP (Figure 3C and 3D). In contrast, the retention of MEX-1(S227A)::GFP in the anterior and posterior is similar to MEX-1::GFP, indicating that S227 is not required for the regulation of MEX-1 retention (Figure 3A, 3B, and 3D). We conclude that PLK-1 phosphorylation of MEX-1 is required to inhibit MEX-1 retention in the anterior cytoplasm, thereby driving MEX-1 segregation to the posterior.

**Figure 3.**
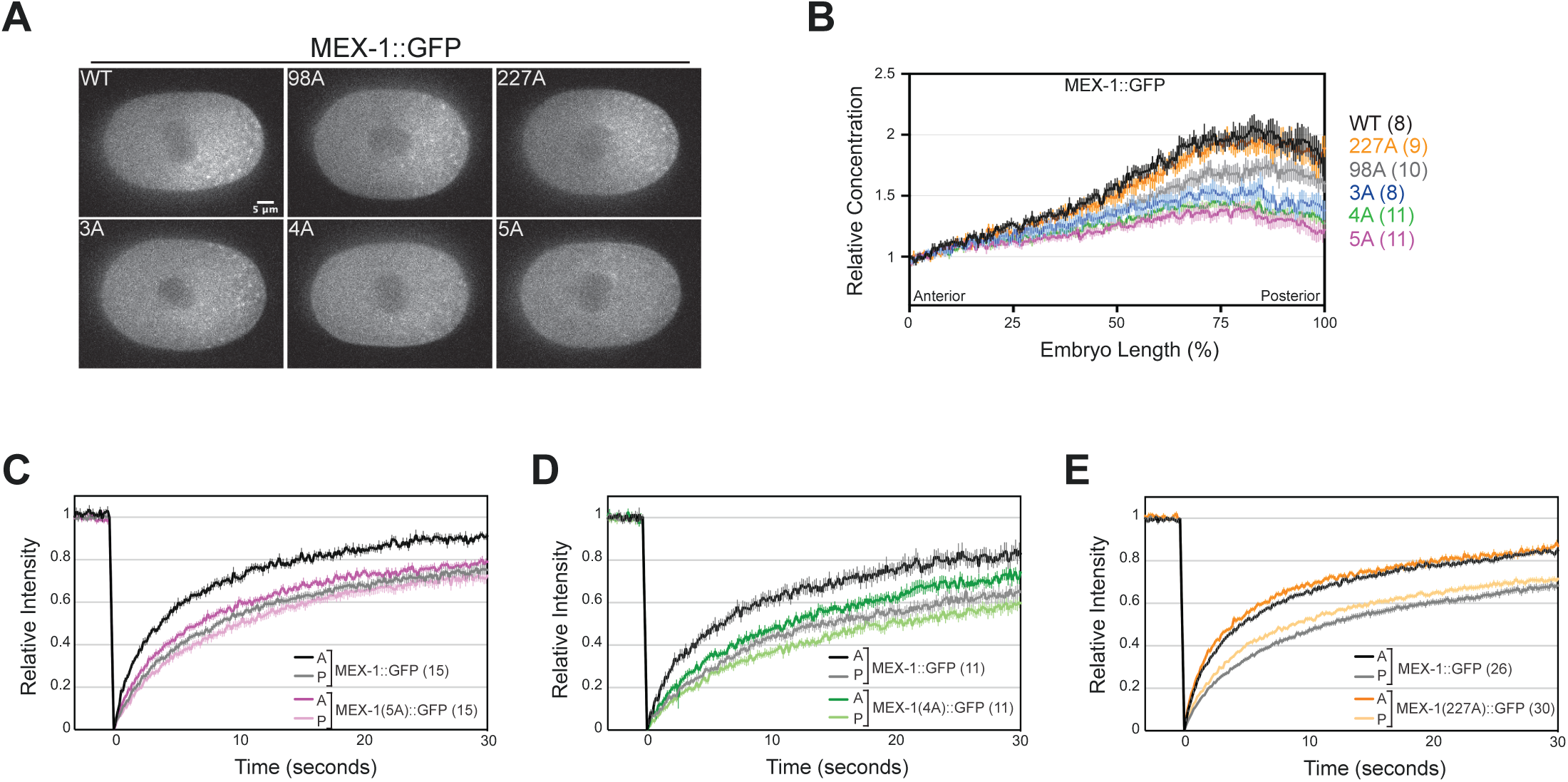
PLK-1 phosphorylation sites are required for MEX-1 segregation. **A.** Fluorescence micrographs of indicated MEX-1::GFP alleles at NEBD. **B.** Average MEX-1::GFP fluorescence intensity along the anteri- or/posterior axis at NEBD. Intensities from the indicated number of embryos were normalized to the anterior end and averaged. **C-E**. FRAP (fluorescence recovery after photobleaching) curves following photobleaching in the anterior (labeled A) or posterior (labeled P) cytoplasm of one-cell embryos at NEBD. The FRAP curves for the indicated number of embryos were normalized and averaged.

### Mutation of MEX-1 phosphorylation sites causes sterility and reduced fecundity at elevated temperatures

The data above suggest that PLK-1 phosphorylation controls MEX-1 segregation similar to its control of POS-1 segregation (Han et al., 2018). To test the functional importance of the PLK-1 phosphorylation of MEX-1 and POS-1, we sought to introduce alanine substitutions at the PLK-1 phosphorylation sites on MEX-1 (S98, T235, S240 and S248) and POS-1 (S199 and S216) (Han et al., 2018). We injected wildtype (N2) worms to avoid potential complications due to epitope tags on POS-1 or MEX-1. Homozygous *mex-1(S98A, T235A, S240A, S248A)* mutants (*mex-1(4A)* hereafter) could be readily maintained at 15°C, 20°C or 25°C and had similar embryonic viability, sterility and brood sizes as wildtype worms at 25°C (Figure 4A – 4C). At the elevated temperature of 25.5°C, *mex-1(4A)* mutants displayed reduced embryonic viability, increased sterility and had smaller brood sizes than wildtype worms (Figure 4A – 4C). In our multiple attempts to isolate *pos-1(S199A;S216A)* mutants, we were only able to independently isolate two *pos-1(S199A;S216A)* heterozygous mutants, both of which gave rise to very few homozygous mutant progeny (3 and 6 progeny total), all of which either died during embryogenesis (2/3 and 5/6 embryos) or were sterile (1/3 and 1/6 adults). As a result, we were unable to maintain either *pos-1(S199A;S216A)* lines. We were able to isolate and maintain homozygous *pos-1(S199A)* mutants, indicating that worms can tolerate mutation of one but not both PLK-1 phosphorylation sites on POS-1. We conclude that PLK-1 phosphorylation is essential for POS-1 function and contributes to MEX-1 function, particularly at elevated temperatures.

**Figure 4.**
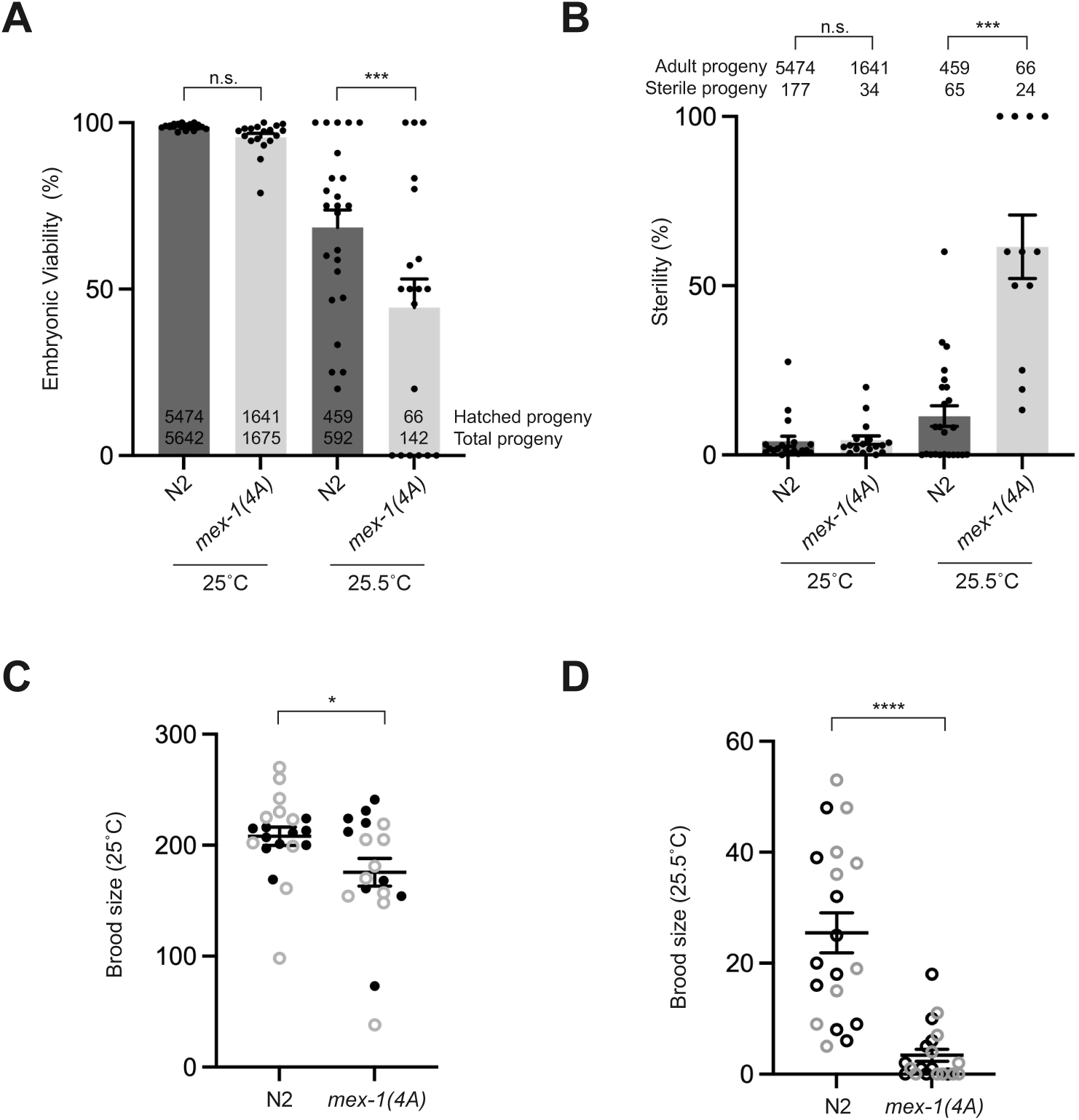
*mex-1(4A)* causes sterility and reduced fecundity at elevated temperatures. **A.** Embryonic viability of N2 (wildtype) and *mex-1(4A)* at the indicated temperatures. Each dot indicates the percentage of viable progeny of an individual hermaphrodite. The total number of hatched embryos (upper) and the total number of embryos laid (lower) is indicated within the graph. **B.** Sterility of N2 and *mex-1(4A)* hermaphrodites at the indicated temperatures. Percentage of sterile progeny from individual hermaphrodites are indicated by black dots. The number of adult and sterile progeny is indicated within the graph. **C,D.** Brood size of N2 and *mex-1(4A)* worms at 25°C and 25.5°C. Each circle indicates the brood size of an individual hermaphrodite. Dark and grey circles indicate technical replicates. Statistical significance for A and B were determined by ANOVA analysis with pairwise post-hoc analysis. Statistical significance for C and D were determined by t-test with Welch’s correction.

In addition to the formation of the posterior-rich MEX-1 gradient, ZIF-1-mediated degradation of MEX-1 in somatic cells contributes to the restriction of MEX-1 to the germline lineage (DeRenzo et al., 2003). We wondered whether the degradation of MEX-1 in somatic cells was important for the viability of *mex-1(4A)* embryos. Depletion of ZIF-1 did not affect the segregation of either MEX-1::GFP or MEX-1(4A)::GFP to the posterior of the zygote, but did prevent their degradation in somatic cells, as expected (Figure 5A-C). At the 4-cell stage, the stabilization of MEX-1 in somatic cells in *zif-1(RNAi)* embryos decreased the ratio of MEX-1::GFP in P2 relative to ABa from 10.8 in control RNAi embryos to 4.0 in *zif-1(RNAi)* embryos and decreased the ratio of MEX-1(4A)::GFP concentration in P2 relative to ABa from 4.4 in control RNAi embryos to 2.7 in *zif-1(RNAi)* embryos (Figure 5D). Nonetheless, *zif-1(RNAi)* did not increase the lethality of either MEX-1::GFP or MEX-1(4A)::GFP embryos (Figure 5E). In addition, we find that reducing the number of P granules by depleting GLH-1, GLH-4, PGL-1 and PGL-3 (“quad RNAi”) (Updike et al., 2014), did not alter the MEX-1::GFP segregation or embryonic viability of MEX-1::GFP or MEX-1(4A)::GFP (Figure 5A-E) embryos.

**Figure 5.**
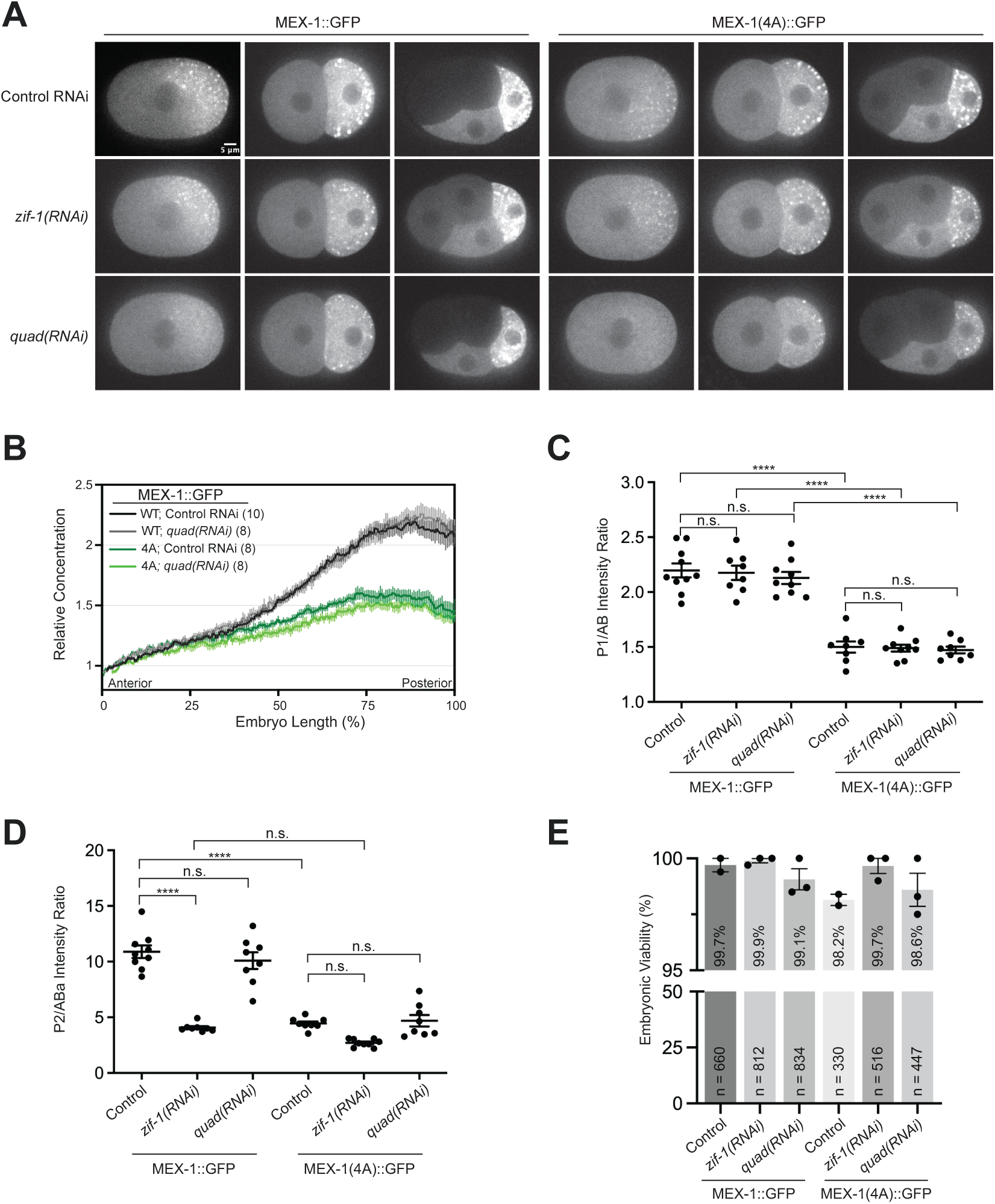
Weakened MEX-1 segregation does not cause embryonic lethality. **A.** Fluorescence micrographs of MEX-1::GFP in 1, 2 and 4-cell embryos following the indicated RNAi depletion. *quad(RNAi)* indicates a combined RNAi targeting the P granule proteins PGL-1, PGL-3, GLH-1 and GLH-4 (Updike et al, 2014). **B.** Average MEX-1::GFP fluorescence intensity along the anterior/posterior axis at NEBD. Intensities from the indicated number of embryos were normalized to the anterior end and averaged. **C.** Ratio of P1/AB fluorescence intensity of MEX-1::GFP in 2-cell embryos. **D.** Ratio of P2/ABa fluorescence intensity of MEX-1::GFP in 4-cell embryos. For C and D, each dot is the ratio in an individual embryo. **E.** Embryonic viability of indicated MEX-1::GFP alleles in the given RNAi condition. The percentage of viable embryos and the total number of embryos analyzed (n) are indicated within the graph. Each black dot indicates the average viability from independent replicates. For all graphs, error bars indicate SEM. Statistical significance in panels C and D was determined by ANOVA analysis with pairwise post-hoc analysis. **** p < 0.0001; n.s. = not significant (> 0.05).

## DISCUSSION

Asymmetric cell division requires coordination between cell cycle and cell polarity mechanisms to ensure that the partitioning of factors is coordinated with cell division. During the asymmetric cell division of *Drosophila* neuroblasts, Polo regulates both PAR polarity through phosphorylation of PAR6 (Wirtz-Peitz et al., 2008) and the segregation of basal determinants through phosphorylation of PON (Wang et al., 2007). During the asymmetric division of the worm zygote, the mitotic kinases PLK-1 and/or Aurora-A coordinate the timing of polarization before and during symmetry breaking (Kapoor and Kotak, 2019; Klinkert et al., 2019; Manzi et al., 2023; Reich et al., 2019; Zhao et al., 2019) and during polarization (Dickinson et al., 2017; Han et al., 2018). In this study, we provide evidence that PLK-1 additionally acts during polarization to control MEX-1 segregation in the *C. elegans* zygote. PLK-1 phosphorylation inhibits MEX-1 retention in the anterior, thereby stimulating MEX-1 accumulation in the posterior cytoplasm. Our findings presented here are similar to our previous findings related to PLK-1 regulation of POS-1 segregation (Han et al., 2018), suggesting similar mechanisms may underlie the segregation of both proteins.

MEX-1 is one of several essential cell fate regulators that localize to a subset of cells during the asymmetric divisions of the early embryo. Although the localization of MEX-1 to posterior cells was significantly weakened by mutation of the PLK-1 phosphorylation sites combined with disruption of P granules or depletion of ZIF-1, we did not observe high levels of embryonic lethality. One possibility is that low levels of MEX-1 asymmetry, for example in *mex-1(4A);zif-1(RNAi)* embryos, is functionally important and that complete disruption of MEX-1 asymmetry would cause embryonic lethality. Alternately, MEX-1 asymmetry may not be required for MEX-1 function. Indeed, the RNA-binding protein MEX-3 retains asymmetric activity even when its asymmetric localization in the early embryo is disrupted (Huang and Hunter, 2015). Additionally, the dramatic enrichment of P granules in the P lineage is not required for the specification of germline (Gallo et al., 2010). In contrast to MEX-1, we were unable to maintain strains in which the PLK-1 phosphorylation sites on POS-1 were mutated, suggesting the asymmetric inheritance of POS-1 may be essential. Taken together, these findings suggest localization to specific lineages may only be important for some fate regulators and highlight the importance of characterizing and disrupting the localization mechanisms of individual proteins in assessing which asymmetries are essential.

Does PLK-1 regulate the segregation of other germplasm components? Both the RNA-binding protein PIE-1 and P granules and are partitioned to the posterior cytoplasm at the same time as MEX-1 and POS-1. The segregation of both PIE-1 and P granules depends on MEX-5/6, PLK-1 and PLK-2 and MBK-2 kinase, which primes the interaction between MEX-5/6 and PLK-1 (Nishi et al., 2008; Pang et al., 2004; Pellettieri et al., 2003; Quintin et al., 2003; Schubert et al., 2000). Because neither MEX-1 nor POS-1 is required for P granule or PIE-1 segregation in the one-cell embryo (Mello et al., 1992; Schnabel et al., 1996; Tabara et al., 1999; Tenenhaus et al., 1998), these observations raise the possibility that phosphorylation by the MEX-5/6/PLK-1 complex may contribute to PIE-1 and/or P granule segregation. Consistent with the possibility that PIE-1 could be a PLK substrate, there are eleven predicted PLK phosphorylation on PIE-1, including two predicted PLK-1 phosphorylation sites (T220 and T308) within the C-terminal region (amino acids 173-335) required for PIE-1 segregation (Reese et al., 2000). However, to our knowledge, phosphorylation of predicted PLK sites on either PIE-1 or P granule proteins has not been reported. Testing whether PLK-1 has a direct role in regulation of PIE-1 and/or P granule segregation will be an interesting avenue for future studies.

## Materials and Methods

### C. elegans strains and maintenance

All strains were derived from the Bristol N2 strain and were maintained on Nematode Growth Medium (NGM) plates containing 3 g/L NaCl, 2.5 g/L peptone and 20 g/L agar supplemented with 1 mM CaCl_2_, 1 mM MgSO_4_, 25 mM KPO_4_ and 5 mg/L Cholesterol with *E. coli* OP50 as a source of food (Brenner, 1974). All RNAi experiments were conducted using the feeding protocol (Timmons and Fire, 1998). *mex-5/6 (RNAi), zif-1(RNAi) and quad(RNAi)* were performed by placing L4 animals on RNAi plates (NGM plates supplemented with 1 mM IPTG and 1.64 mM carbenicillin) for 24 hours at room temperature. Strains used in this study are listed in Table 3.

### Gene editing

CRISPR/Cas9 gene editing was performed similar to the method described in Ghanta et al., 2021 (Ghanta et al., 2021). Briefly, injection mixtures containing 400 ng pRF4::*rol-6(su1006)* plasmid, 30 pmol Cas9 (IDT, Cat#1081058), 90 pmol tracrRNA (IDT, Cat#1072532) 95 pmol crRNA, 1100 ng ssDNA oligo donor and nuclease free water (Bio Basic) were injected into L4 hermaphrodites. Injected worms were singled onto individual plates. F1 Rollers were singled, allowed to lay embryos, and genotyped by PCR. Homozygous F2s animals were identified by PCR genotyping and validated by sequencing. pRF4::*rol-6(su1006)* plasmid was purified using PureLink HiPure plasmid miniprep kit (Invitrogen, Cat#K210003). Single worm PCR was performed by picking individual worms into PCR tubes containing 1X Taq reaction buffer (NEB, Cat#B9014S), Proteinase K (Roche; Cat#03115828001) and nuclease free water, freezing at -80°C for at least 15 min, and lysing at 60°C for 1 hr, following by 95°C for 15 min. PCR reaction mixes containing 1X *Taq* reaction buffer, 10 µM primers (IDT), 10 mM dNTPs (NEB; Cat#N0447S), Taq DNA polymerase (NEB; Cat#M0273X) and nuclease free water were added to 5 µL lysate. DNA oligonucleotides and cRNAs used for gene editing are listed in Tables 1 and 2.

**Table 1.**
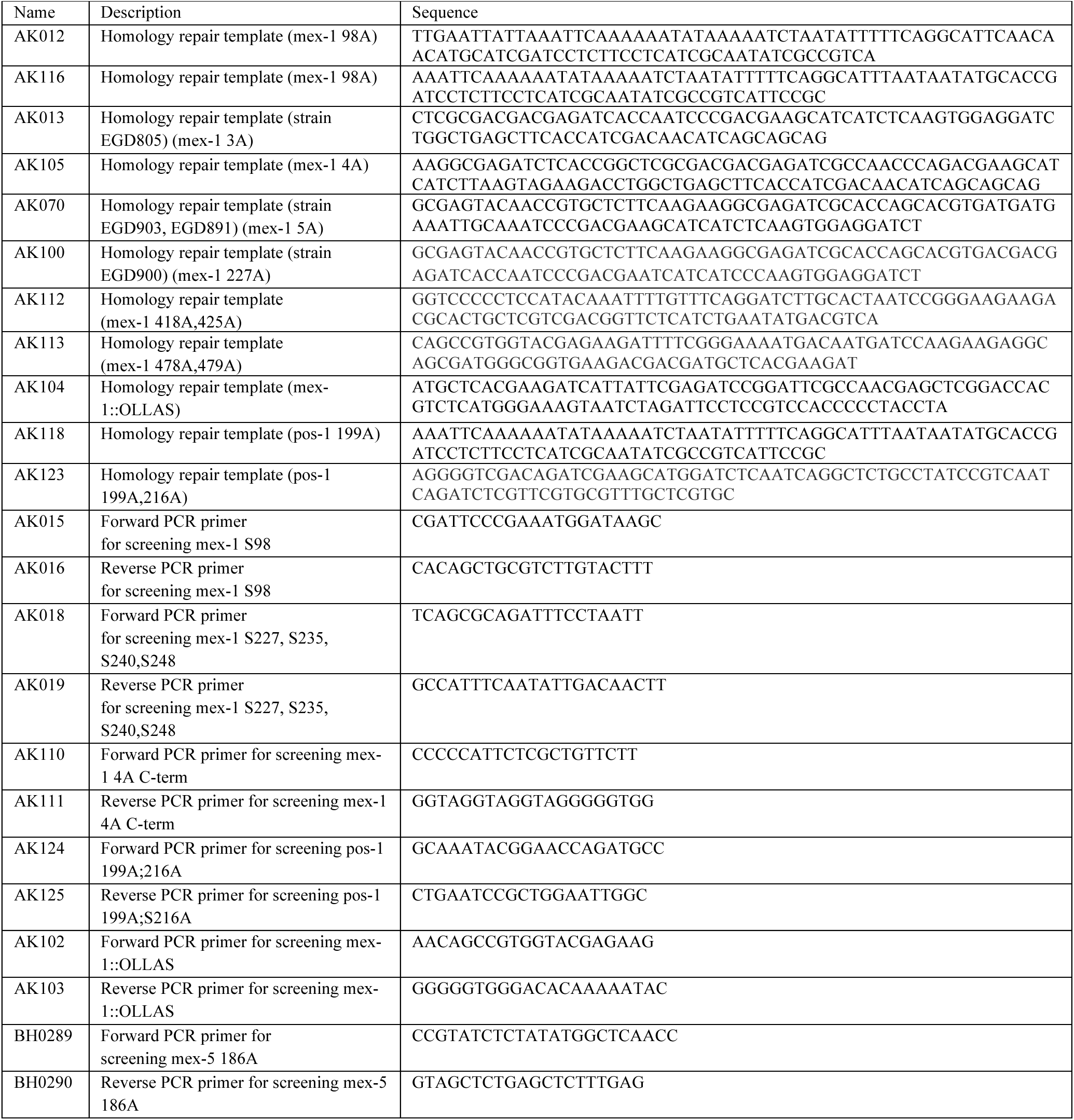
DNA oligonucleotides used in this study.

**Table 2.**
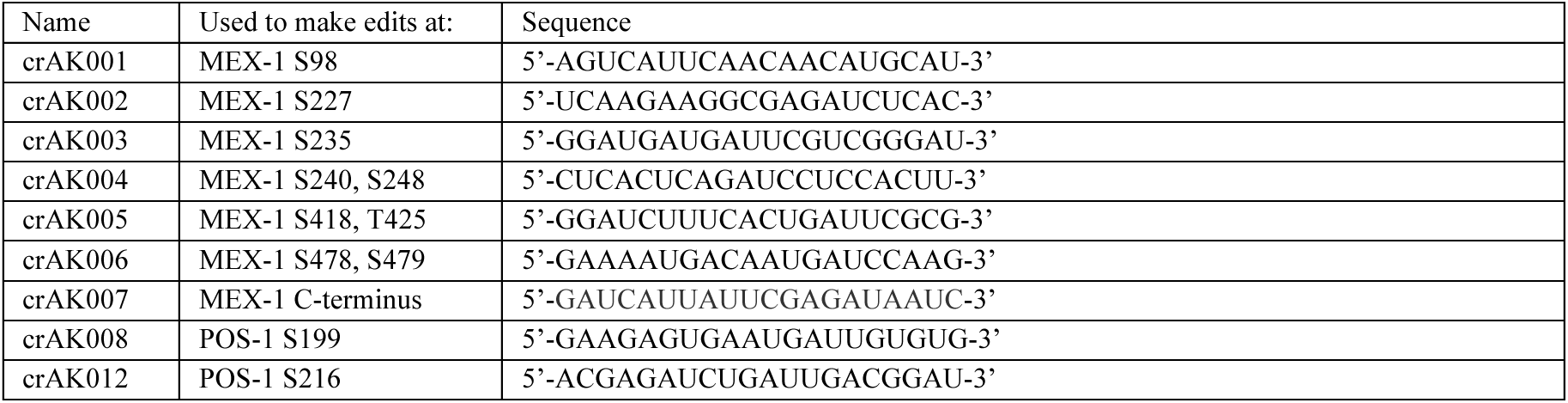
crRNAs used in this study.

**Table 3.** Strains used in this study.

| Strain | Genotype | Construction | Reference: |
| --- | --- | --- | --- |
| EGD135<br>mex-1::gfp | mex-1(egx9[mex-1::gfp]) II |  | Gauvin (2018) |
| EGD799<br>mex-1(S98A) | mex-1(egx116[mex-1(S98A)]) II | HR template: AK116<br>crRNA: crAK001<br>Injected N2 | This study |
| EGD800<br>mex-1(S98A)::gfp | mex-1(egx117[mex-1(S98A)::gfp]) II | HR template: AK012<br>crRNA: crAK001<br>Injected EGD135 | This study |
| EGD805<br>mex-1(3A)::gfp | mex-1(egx122[mex-1(S98A, S240A, S248A)::gfp]) | HR template: AK071<br>crRNA: crAK004<br>Injected EGD799 | This study |
| EGD1012<br>mex-1(4A)::gfp | mex-1(egx242[mex-1(S98A, T235A, S240A, S248A)::gfp]) II | HR template: AK013<br>crRNA: crAK004<br>Injected EGD800 | This study |
| EGD1013<br>mex-1(4A) | mex-1(egx243[mex-1(S98A, T235A, S240A, S248A)]) II | HR template: AK105<br>crRNA: crAK003,<br>crAK004<br>Injected EGD799 | This study |
| EGD903<br>mex-1(5A)::gfp | mex-1(egx124[mex-1 (S98A, S227A, S235A, S240A, S248A)::gfp]) II | HR template: AK070<br>crRNA: crAK002,<br>crAK003<br>Injected EGD805 | This study |
| EGD891<br>mex-1(5A) | mex-1(egx181[mex-1(S98A, S227A, S235A, S240A, S248A)]) II | HR template: AK070<br>crRNA: crAK002,<br>crAK003<br>Injected EGD889 | This study |
| EGD906<br>mex-1(5A)::OLLAS | mex-1(egx190[mex-1::ollas(S98A, S227A, T235A, S240A, S248A)]) II | HR template: AK104<br>crRNA: crAK007<br>Injected EGD886 | This study |
| EGD900<br>mex-1(S277A)::gfp | mex-1(egx187[mex-1::gfp(S277A)]) II | HR template: AK100<br>crRNA: crAK002<br>Injected EGD135 | This study |
| EGD913<br>mex-1::ollas | mex-1(egx197[mex-1::ollas]) II | HR template: AK104<br>crRNA: crAK007<br>Injected N2 | This study |
| EGD1011 | mex-1(egx9[mex-1::gfp]) II; (mex-5(T186A)) V | Crossed EGD135 to EGD298 | This study |
| EGD1017<br>mex-1(4A-C) | mex-1(egx247[mex-1(S418A, S425A, S478A, S479A)]) II | Rnd 1:HR: AK113,<br>crRNA: crAK006<br>Rnd2: HR: AK112<br>crRNA: crAK005<br>Injected N2 | This study |
| EGD1019<br>mex-1(4A-C)::ollas | mex-1(egx247[mex-1(S418A, S425A, S478A, S479A)::ollas]) II | HR template AK104<br>crRNA: AK007<br>Injected EGD1017 | This study |
| EGD921 | pos-1(egx202[pos-1(S199A)]) V | HR template: AK118<br>crRNA: crAK008<br>Injected N2 | This study |

### BI2536 treatment

For small molecule inhibitor treatment, embryo eggshells were permeabilized using *perm-1* RNAi (Carvalho et al., 2011). A perm-1(RNAi) bacterial culture was diluted 1:4 with L4440 bacterial culture and seeded on a RNAi plate overnight. MEX-1::GFP L4 animals were fed on RNAi for 18 hours. Permeabilization of the eggshell by *perm-1* RNAi was confirmed using BioTracker 640 Red C2 (FM4-64) dye (Sigma-Aldrich). Embryos were hand-dissected in simple embryonic culture buffer (0.5 μg/μL Inulin, 50mM pH 7.4 HEPES, 20% FCS, 50% L-15 medium) containing 20μm-diameter polystyrene microspheres (Bangs Laboratories, Inc. Cat# NT30N), 33 µM BioTracker 640 Red C2 (FM4-64) dye and either DMSO (Control) or 20 µM BI2536 small molecule inhibitor (Axon MedChem). Embryos were incubated for 5-7 minutes before imaging.

### Protein purification

DNA encoding amino acids 1-299 of MEX-1 and MEX-1(5A) were synthesized (IDT gBlocks), cloned into the protein expression vector pHMTc (Ryder et al., 2004), and transformed into *E.coli* strain BL21(DE3). Bacteria were grown in TB buffer and induced with 0.5mM IPTG and cultured overnight at 16°C. 100 µM Zn(OAc)_2_ was added at the time of induction. Bacterial pellets were lysed in 50 mM Tris-HCl, pH 8.0, 1 M NaCl, 20 mM imidazole, 5 mM BME, and cOmplete EDTA-free Protease Inhibitor cocktail (Sigma-Aldrich) using a microfluidizer and clarified by centrifugation at 3000 rpm for 20 min at 4°C. Lysates were bound in batch to Ni-NTA beads (G biosciences), washed in 4 column volumes with wash buffer (50 mM Tris-HCl pH 8.0. 250 mM NaCl, 100 µM Zn(OAc)_2_, 5 mM BME and 20 mM imidazole) and eluted in elution buffer (wash buffer containing 300 mM imidazole). Elution fractions containing MBP:MEX-1(1-299):6XHis were bound in batch to amylose beads (NEB; Cat#E8021S), washed in 4 column volumes with wash buffer (50 mM Tris-HCl pH 8.0, 250 mM NaCl, 100 µM Zn(OAc)_2_, 5 mM BME) and eluted in wash buffer containing 10 mM maltose. Elution fractions containing MBP:MEX-1(1-299):6XHis were pooled, aliquoted, frozen and used for *in vitro* kinase assays.

### In vitro kinase assay and mass spectrometry

Kinase assays were performed at 37°C by diluting 0.2ug of substrate protein into kinase reaction buffer (8 mM MOPS, pH 7.0, 100 μM Zn(OAc)_2_, 10 mM MgCl_2_ and protease cocktail (K1010, APExBIO, Houston, TX, USA) containing 2 mM ATP-γS (Abcam) and 375 ng of hPLK1 (EMD Millipore). Samples were collected at the at the indicated time points and quenched with 20 mM EDTA. To alkylate ATP-γS, final concentration of 2.5 mM P-nitrobenzyl mesylate (Abcam) was added to samples and incubated for 2 hours at room temperature. Kinase reactions were analyzed by Western Blot using 1:5000 thiophosphate ester specific primary antibody (Abcam) and 1:10,000 peroxidase-conjugated AffiniPure Goat anti-rabbit IgG secondary antibody (Jackson ImmunoResearch). Blots were developed with the Clarity Western ECL Substrate (Bio-Rad) and imaged with the ChemiDoc XRS system (Bio-Rad) with auto exposure setting. Reaction samples were also run on SDS-PAGE gels and stained with Coomassie Brilliant Blue to quantify substrate levels.

For phopsho-mass spectrometry analysis, kinase assays were performed as described above except that protease cocktail, 2 mM ATP-γS and P-nitrobenzyl mesylate are not added in the reaction buffer. Samples were run on SDS-PAGE gels and stained with Coommasie Brilliant Blue in a clean petri dish. MEX-1 was excised from the gel and stored in sterile water. For mass spectrometry analysis, gel slices were destained, digested with trypsin, and peptides extracted. Peptides were analyzed on an Orbitrap Fusion Lumos mass spectrometer (Thermo Fisher Scientific) equipped with an Easy-nLC (Thermo Fisher Scientific). Raw data were searched using Comet (Eng et al., 2013) against a custom database containing the MEX1 and MEX1-5A sequences and phosphorylation on S/T/Y as a dynamic modification.

### Embryonic viability, brood size and sterility assay

Figure 4: MEX-1(4A) and N2 worms were passaged and synchronized at 25°C prior to the experiment. For the experiments performed at 25.5°C, L4 worms at 25°C were transferred to fresh plates and incubated at 25°C or 25.5°C and allowed to lay progeny. When F1s reached the L4 stage, 10 individual worms were singled to NGM plates (IPM; Cat#11006-518) seeded with OP50 and incubated at the indicated temperatures. The animals were transferred to fresh plates every 24 hours until the animals died. Embryos laid and the number of surviving L1s were. F1 animals that did not lay embryos were counted as sterile. On plates with non-sterile F1s, the number of F2 embryos and surviving L1s were counted to determine embryonic lethality.

Figure 5: To determine the embryonic viabilityMEX-1::GFP and MEX-1(4A)::GFP following RNAi, L4 worms were transferred to *quad*(RNAi) and *zif-1*(RNAi) plates and incubated for 24 hours at 23°C. 4 young adult animals were moved to 20°C and allowed to lay embryos on fresh RNAi plates for three 2-hour intervals. and then transferred to fresh corresponding RNAi plates for 6 hours at 20°C. Embryonic viability was calculated as the percentage of embryos laid that hatched.

### Microscopy

Images were collected on a Marianas spinning disk confocal microscope controlled by the Slidebook software package (Intelligent Imaging Innovations, Denver, CO) and built around a Zeiss Axio Observer Z.1 equipped with a Zeiss Plan-Apochromat 63×/1.4NA oil immersion objective, a CSU-X1 spinning disk (Yokogawa, Tokyo, Japan), an Evolve 512X512 EMCCD camera (Photometrics, Tucson, AZ) and a 50mW 488nm solid state laser.

FRAP experiments were performed using a Phasor photomanipulation unit (Intelligent Imaging Innovations), which delivered 488nm light simultaneously to two 5 μm diameter circular ROIs, one in the anterior and one in the posterior cytoplasm during NEBD in one-cell embryo. Photobleaching lasted for 90 msec and images were collected at 93.1 msec per frame for 300 frames.

### Quantification and statistical analysis

To quantify *in vitro* kinase assays, pixel intensities of blot images were inverted using ImageJ. Identically sized regions of interest (ROIs) were used to measure the pixel intensities of the protein bands and background. Net values of protein bands were defined by deducting the inverted background from the inverted band value. The final relative quantification values are defined as the ratio of a net band value to the final MBP:MEX-1(aa1-299):His time point value.

## Acknowledgements

We thank members of the Griffin lab, Jamie Moseley, Monica Gotta and Amanda Amodeo for helpful discussions. We thank Ann Lavanway for her support.

---

**Supplemental Figure 1.**
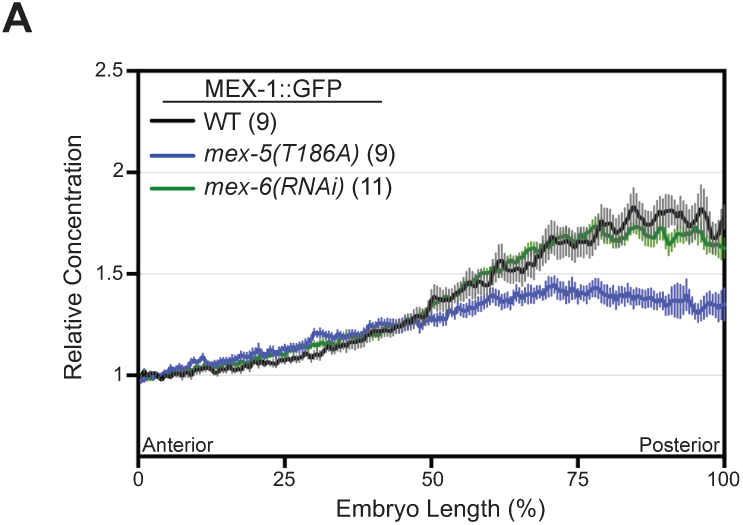
Average MEX-1::GFP fluorescence intensity along the anterior/posterior axis *mex-5(T186A)* and *mex-6(RNAi)* embryos at NEBD. Intensities from the indicated number of embryos were normalized to the anterior end and averaged.

## References

Barbieri, S., A. Nurni Ravi, E.E. Griffin, and M. Gotta. 2022. Modeling protein dynamics in *Caenorhabditis elegans* embryos reveals that the PLK-1 gradient relies on weakly coupled reaction-diffusion mechanisms. Proc Natl Acad Sci U S A. 119:e2114205119.

Bowerman, B., B.W. Draper, C.C. Mello, and J.R. Priess. 1993. The maternal gene skn-1 encodes a protein that is distributed unequally in early *C. elegans* embryos. Cell. 74:443–452.

Brenner, S. 1974. The genetics of Caenorhabditis elegans. Genetics. 77:71–94.

Budirahardja, Y., and P. Gonczy. 2008. PLK-1 asymmetry contributes to asynchronous cell division of *C. elegans* embryos. Development. 135: 1303–1313.

Carvalho, A., S.K. Olson, E. Gutierrez, K. Zhang, L.B. Noble, E. Zanin, A. Desai, A. Groisman, and K. Oegema. 2011. Acute drug treatment in the early *C. elegans* embryo. PLoS ONE. 6:e24656–e24656.

Chase, D., C. Serafinas, N. Ashcroa, M. Kosinski, D. Longo, D.K. Ferris, and A. Golden. 2000. The polo-like kinase PLK-1 is required for nuclear envelope breakdown and the completion of meiosis in *Caenorhabditis elegans*. Genesis. 26:26–41.

Daniels, B.R., T.M. Dobrowsky, E.M. Perkins, S.X. Sun, and D. Wirtz. 2010. MEX-5 enrichment in the *C. elegans* early embryo mediated by differential diffusion. Development. 137:2579–2585.

Daniels, B.R., E.M. Perkins, T.M. Dobrowsky, S.X. Sun, and D. Wirtz. 2009. Asymmetric enrichment of PIE-1 in the *Caenorhabditis elegans* zygote mediated by binary counterdiffusion. J Cell Biol. 184:473–479.

Detienzo, C., K.J. Reese, and G. Seydoux. 2003. Exclusion of germ plasm proteins from somatic lineages by cullin-dependent degradation. Nature. 424:685–689.

Dickinson, D.J., F. Schwager, L. Pintard, M. Gotta, and B. Goldstein. 2017. A Single-Cell Biochemistry Approach Reveals PAR Complex Dynamics during Cell Polarization. Dev Cell. 42:416–434 e411.

Elia, A.E., P. Rellos, L.F. Haire, J.W. Chao, F.J. Ivins, K. Hoepker, D. Mohammad, L.C. Cantley, S.J. Smerdon, and M.B. Yaffe. 2003. The molecular basis for phosphodependent substrate targeting and regulation of Plks by the Polo-box domain. Cell. 115:83–95.

Eng, J.K., T.A. Jahan, and M.R. Hoopmann. 2013. Comet: an open-source MS/MS sequence database search tool. Proteomics. 13:22–24.

Gallo, C.M., J.T. Wang, F. Motegi, and G. Seydoux. 2010. Cytoplasmic parJJoning of P granule components is not required to specify the germline in *C. elegans*. Science. 330:1685–1689.

Gauvin, T.J., B. Han, M.J. Sun, and E.E. Griffin. 2018. PIE-1 translation in the germline lineage contributes to PIE-1 asymmetry in the early *Caenorhabditis elegans* embryo. G3. 8:3791–3801.

Ghanta, K.S., T. Ishidate, and C.C. Mello. 2021. Microinjection for precision genome editing in *Caenorhabditis elegans*. STAR Protoc. 2:100748.

Griffin, E.E. 2015. Cytoplasmic localization and asymmetric division in the early embryo of *Caenorhabditis elegans*. Wiley Interdiscip Rev Dev Biol. 4:267–282.

Griffin, E.E., D.J. Odde, and G. Seydoux. 2011. Regulation of the MEX-5 gradient by a spatially segregated kinase/phosphatase cycle. Cell. 146:955–968.

Guedes, S., and J.R. Priess. 1997. The *C. elegans* MEX-1 protein is present in germline blastomeres and is a P granule component. Development. 124:731–739.

Han, B., K.R. Antkowiak, X. Fan, M. Rutigliano, S.P. Ryder, and E.E. Griffin. 2018. Polo-like Kinase couples cytoplasmic protein gradients in the *C. elegans* zygote. Curr Biol. 28:60–69.

Holzer, E., C. Rumpf-Kienzl, S. Falk, and A. Dammermann. 2022. A modified TurboID approach idenJfies Jssue-specific centriolar components in *C. elegans*. PLoS Genet. 18:e1010150.

Huang, N.N., and C.P. Hunter. 2015. The RNA binding protein MEX-3 retains asymmetric activity in the early *Caenorhabditis elegans* embryo in the absence of asymmetric protein localization. Gene. 554:160–173.

Hwang, S.-Y., and L.S. Rose. 2010. Control of asymmetric cell division in early *C. elegans* embryogenesis: teaming-up translational repression and protein degradation. BMB Rep. 43:69–78.

Kapoor, S., and S. Kotak. 2019. Centrosome Aurora A regulates RhoGEF ECT-2 localisation and ensures a single PAR-2 polarity axis in *C. elegans* embryos. Development. 146:dev174565.

Klinkert, K., N. Levernier, P. Gross, C. Gentili, L. von Tobel, M. Pierron, C. Busso, S. Herrman, S.W. Grill, K. Kruse, and P. Gonczy. 2019. Aurora A depletion reveals centrosome-independent polarization mechanism in *Caenorhabditis elegans*. Elife. 8.:e44552.

Kumar, M., S. Michael, J. Alvarado-Valverde, B. Mészáros, H. Sámano-Sánchez, A. Zeke, L. Dobson, T. Lazar, M. Örd, A. Nagpal, N. Farahi, M. Käser, R. KraleJ, N.E. Davey, R. Pancsa, L.B. Chemes, and T.J. Gibson. 2022. The Eukaryotic Linear Motif resource: 2022 release. Nucleic Acids Res. 50:D497–D508.

Lang, C.F., and E. Munro. 2017. The PAR proteins: from molecular circuits to dynamic self-stabilizing cell polarity. Development. 144:3405–3416.

Li, R. 2013. The art of choreographing asymmetric cell division. Dev Cell. 25:439–450.

Liu, Z., J. Ren, J. Cao, J. He, X. Yao, C. Jin, and Y. Xue. 2013. Systematic analysis of the Plk-mediated phosphoregulation in eukaryotes. Brief Bioinform. 14:344–360.

Manzi, N.I., B.N. de Jesus, Y. Shi, and D.J. Dickinson. 2024. Temporally distinct roles of Aurora A in polarization of the *C. elegans* zygote. Development. 151:dev202479.

Mello, C.C., B.W. Draper, M. Krause, H. Weintraub, and J.R. Priess. 1992. The pie-1 and mex-1 genes and maternal control of blastomere identity in early *C. elegans* embryos. Cell. 70: 163–176.

Mello, C.C., C. Schubert, B. Draper, W. Zhang, R. Lobel, and J.R. Priess. 1996. The PIE-1 protein and germline specification in *C. elegans* embryos. Nature. 382:710–712.

Moreno, S., and P. Nurse. 1990. Substrates for p34cdc2: in vivo veritas? Cell. 61:549–551.

Nakajima, H., F. Toyoshima-Morimoto, E. Taniguchi, and E. Nishida. 2003. IdenJfication of a consensus motif for Plk (Polo-like kinase) phosphorylation reveals Myt1 as a Plk1 substrate. J Biol Chem. 278:25277–25280.

Nigg, E.A. 1993. Cellular substrates of p34cdc2 and its companion cyclin-dependent kinases. Trends in Cell Biology. 3:296–301.

Nishi, Y., E. Rogers, S.M. Robertson, and R. Lin. 2008. Polo kinases regulate *C. elegans* embryonic polarity via binding to DYRK2-primed MEX-5 and MEX-6. Development. 135:687–697.

Offenburger, S.L., D. Bensaddek, A.B. Murillo, A.I. Lamond, and A. Gartner. 2017. Comparative genetic, proteomic and phosphoproteomic analysis of *C. elegans* embryos with a focus on ham-1/STOX and pig-1/MELK in dopaminergic neuron development. Sci Rep. 7:4314.

Oldenbroek, M., S.M. Robertson, T. Guven-Ozkan, S. Gore, Y. Nishi, and R. Lin. 2012. Multiple RNA-binding proteins function combinatorially to control the soma-restricted expression pattern of the E3 ligase subunit ZIF-1. Dev Biol. 363:388–398.

Oldenbroek, M., S.M. Robertson, T. Guven-Ozkan, C. Spike, D. Greenstein, and R. Lin. 2013. Regulation of maternal Wnt mRNA translation in *C. elegans* embryos. Development. 140:4614–4623.

Pang, K.M., T. Ishidate, K. Nakamura, M. Shirayama, C. Trzepacz, C.M. Schubert, J.R. Priess, and C.C. Mello. 2004. The minibrain kinase homolog, mbk-2, is required for spindle positioning and asymmetric cell division in early *C. elegans* embryos. Dev Biol. 265:127–139.

Peglion, F., and N.W. Goehring. 2019. Switching states: dynamic remodelling of polarity complexes as a toolkit for cell polarization. Curr Opin Cell Biol. 60:121–130.

Pelleperi, J., V. Reinke, S.K. Kim, and G. Seydoux. 2003. Coordinate activation of maternal protein degradation during the egg-to-embryo transition in *C. elegans*. Dev Cell. 5:451–462.

Quintin, S., P.E. Mains, A. Zinke, and A.A. Hyman. 2003. The mbk-2 kinase is required for inactivation of MEI-1/katanin in the one-cell *Caenorhabditis elegans* embryo. EMBO Rep. 4:1175–1181.

Reese, K.J., M.A. Dunn, J.A. Waddle, and G. Seydoux. 2000. Asymmetric segregation of PIE-1 in *C. elegans* is mediated by two complementary mechanisms that act through separate PIE-1 protein domains. Mol Cell. 6:445–455.

Reich, J.D., L. Hubatsch, R. Illukkumbura, F. Peglion, T. Bland, N. Hirani, and N.W. Goehring. 2019. Regulated Activation of the PAR Polarity Network Ensures a Timely and Specific Response to Spatial Cues. Curr Biol. 29:1911–1923.

Rivers, D.M., S. Moreno, M. Abraham, and J. Ahringer. 2008. PAR proteins direct asymmetry of the cell cycle regulators Polo-like kinase and Cdc25. J Cell Biol. 180:877–885.

Rose, L., and P. Gonczy. 2014. Polarity establishment, asymmetric division and segregation of fate determinants in early *C. elegans* embryos. Wormtiook 1–43.

Ryder, S.P., L.A. Frater, D.L. Abramovitz, E.B. Goodwin and J.R. Williamson. 2004. RNA target specificity of the STAR/GSG domain post-transcriptional regulatory protein GLD-1. Nat Struc Mol Biol. 11:20–28.

Schnabel, R., C. Weigner, H. Hutter, R. Feichtinger, and H. Schnabel. 1996. mex-1 and the general parJJoning of cell fate in the early *C. elegans* embryo. Mechanisms of Development. 54:133–147.

Schubert, C.M., R. Lin, C.J. de Vries, R.H. Plasterk, and J.R. Priess. 2000. MEX-5 and MEX-6 function to establish soma/germline asymmetry in early *C. elegans* embryos. Mol Cell. 5:671–682.

Songyang, Z., S. Blechner, N. Hoagland, M.F. Hoekstra, H. Piwnica-Worms, and L.C. Cantley. 1994. Use of an oriented peptide library to determine the optimal substrates of protein kinases. Current Biology. 4:973–982.

Steegmaier, M., M. Hoffmann, A. Baum, P. Lenart, M. Petronczki, M. Krssak, U. Gurtler, P. Garin-Chesa, S. Lieb, J. Quant, M. Grauert, G.R. Adolf, N. Kraut, J.M. Peters, and W.J. Rettig. 2007. BI2536, a potent and selective inhibitor of polo-like kinase 1, inhibits tumor growth in vivo. Curr Biol. 17:316–322.

Sunchu, B., and C. Cabernard. 2020. Principles and mechanisms of asymmetric cell division. Development. 147:dev167650.

Tabara, H., R.J. Hill, C.C. Mello, J.R. Priess, and Y. Kohara. 1999. pos-1 encodes a cytoplasmic zinc-finger protein essential for germline specification in *C. elegans*. Development. 126:1–11.

Tenenhaus, C., C. Schubert, and G. Seydoux. 1998. Genetic requirements for PIE-1 localization and inhibition of gene expression in the embryonic germ lineage of *Caenorhabditis elegans*. Dev Biol. 200:212–224.

Tenlen, J.R., J.N. Molk, N. London, B.D. Page, and J.R. Priess. 2008. MEX-5 asymmetry in one-cell *C. elegans* embryos requires PAR-4- and PAR-1-dependent phosphorylation. Development. 135:3665–3675.

Timmons, L., and A. Fire. 1998. Specific interference by ingested dsRNA. Nature. 395:854.

Updike, D.L., A.K.a. Knutson, T.A. Egelhofer, A.C. Campbell, and S. Strome. 2014. Germ-granule components prevent somatic development in the *C. elegans* germline. Curr Biol. 24:970–975.

Wang, H., Y. Ouyang, W.G. Somers, W. Chia, and B. Lu. 2007. Polo inhibits progenitor self-renewal and regulates Numb asymmetry by phosphorylating Pon. Nature. 449:96–100.

Wang, J.T., and G. Seydoux. 2013. Germ cell specification. Adv. Exp. Med. Biol. 757:17–39.

Wirtz-Peitz, F., T. Nishimura, and J. Knoblich. 2008. Linking Cell Cycle to Asymmetric Division: Aurora-A Phosphorylates the Par Complex to Regulate Numb Localization. Cell. 135:161–173.

Wu, Y., B. Han, Y. Li, E. Munro, D.J. Odde, and E.E. Griffin. 2018. Rapid diffusion-state switching underlies stable cytoplasmic gradients in the *Caenorhabditis elegans* zygote. Proc Natl Acad Sci U S A. 115:E8440–E8449.

Wu, Y., H. Zhang, and E.E. Griffin. 2015. Coupling between cytoplasmic concentration gradients through local control of protein mobility in the *Caenorhabditis elegans* zygote. Mol Biol Cell. 26:2963–2970.

Zhao, P., X. Teng, S.N. Tantirimudalige, M. Nishikawa, T. Wohland, Y. Toyama, and F. Motegi. 2019. Aurora-A breaks symmetry in contractile actomyosin networks independently of its role in centrosome maturation. Dev Cell. 48:631–645.

